# *Cis*-regulatory evolution that caused change in *wingless* expression pattern associated with wing pigmentation pattern of *Drosophila*

**DOI:** 10.1101/2023.03.01.530703

**Authors:** Takumi Karasawa, Namiho Saito, Shigeyuki Koshikawa

## Abstract

Genetic mechanisms underlying the acquisition of new traits are an important topic in evolutionary developmental biology. Especially, the co-option of important regulatory genes potentially plays an important role in the gain of new traits. However, how the co-option occurs at the sequence level is still elusive. *Drosophila guttifera* has a unique wing pigmentation pattern and this is newly gained via the evolution of the expression pattern of *wingless*, which induces the pigmentation pattern formation. In this study, to reveal the changes in the *cis*-regulatory sequence which caused the co-option of *wingless* that lead to the expression in a new place, we conducted transgenic EGFP reporter assays of altered *cis*-regulatory sequences. As a result, the sequence was divided into regions needed to activate expression in the entire wing veins and a region required for repressing expression in excess parts. Comparisons with the homologous sequence of *Drosophila melanogaster* showed that the repressive function of the *cis*-regulatory region is also possessed by *D. melanogaster* while the activating function is newly gained in a lineage leading to *D. guttifera*. Furthermore, a putative binding site of SMAD transcription factors is shown to be essential for activating expression but also existing in the homologous region of *D. melanogaster*. Our results suggest that the pre-existing regulatory sequences in the *cis*-regulatory region coordinate with the newly gained sequences to acquire the new expression pattern of *wingless*.

**Graphical Abstract:** 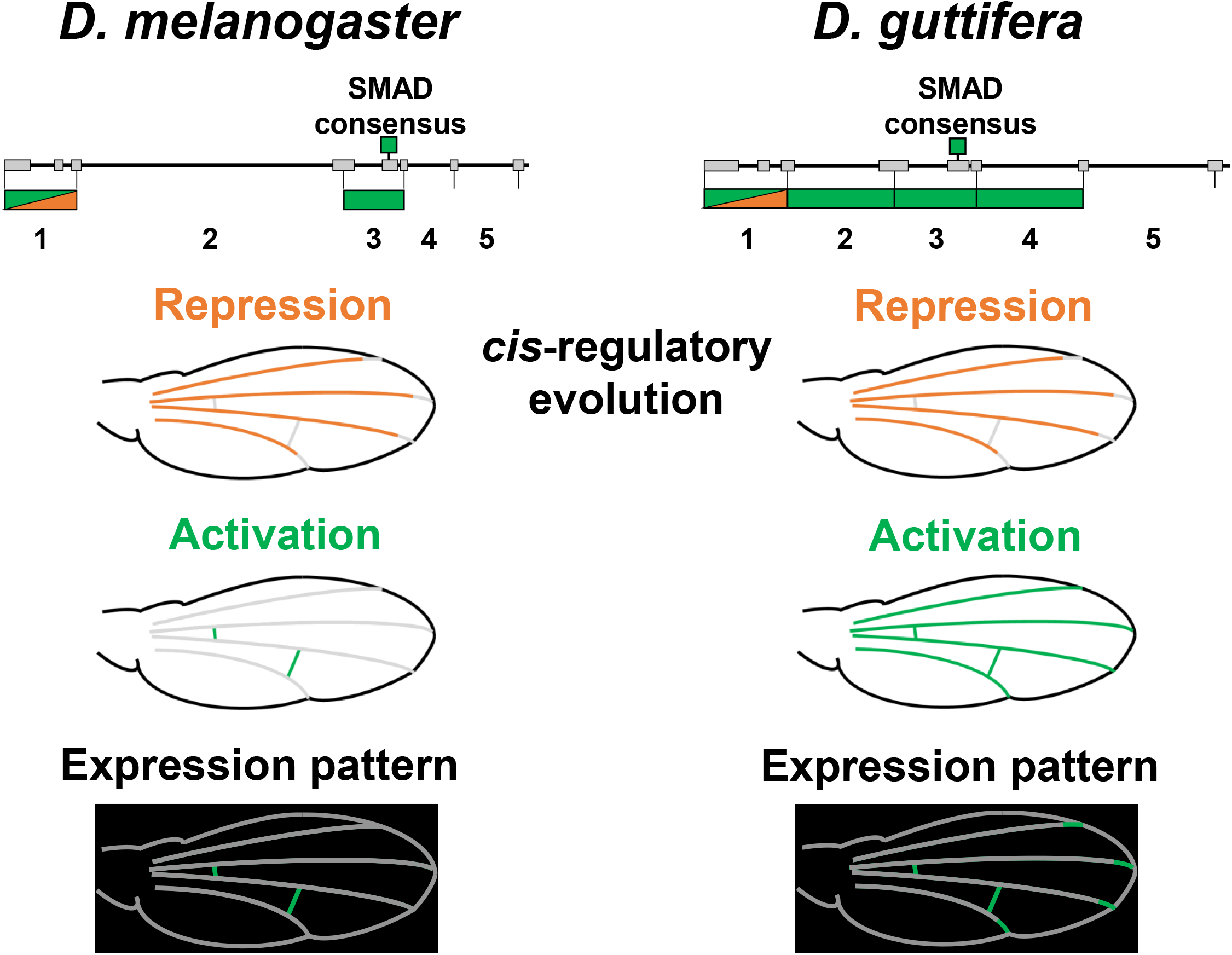

## Introduction

Evolution has produced various traits throughout the history of life. In evolutionary developmental biology, genetic mechanisms underlying the gain of new traits have been discussed [1–3]. In particular, the importance of gene network co-option has been examined [4, 5]. In gene network co-option, a regulatory factor is redeployed in a novel cellular context, and genes in the network controlled by this regulatory factor are also made to function in the context. Thus, changes to a few genes potentially alter the expression of many downstream genes rapidly. There are some examples in which gene network co-option may have played an important role in the gain of new morphological traits [6–10]. However, we do not have a good understanding of how the co-option that leads to the acquisition of traits occurs at the sequence level.

The wing pigmentation pattern of *Drosophila* species is one of the traits that have been studied to reveal the genetic changes underlying the acquisition of traits [11–13]. For example, in *Drosophila biarmipes*, the co-option of *Distal-less (Dll)*, a gene that also functions in the patterning of the wing margin, contributes to the gain of the male-specific spot on the wing [14]. *Drosophila guttifera* has a unique pigmentation pattern on its wing, and the pigmentation is regulated by a gene called *wingless (wg)*, which activates the expression of pigmentation genes such as *yellow* (Fig. 1A) [15, 16]. In *D. guttifera*, *wg* is expressed in the crossveins, longitudinal vein tips, and campaniform sensilla of the pupal wing, prefiguring the location of pigmentation [17, 18]. In *Drosophila melanogaster*, on the other hand, *wg* is expressed in the crossveins, but not in the longitudinal vein tips or campaniform sensilla. Furthermore, in fruit flies of the quinaria group, which is closely related to *D. guttifera* [19], the expression pattern of *wg* is similar to that of *D. melanogaster*, suggesting that the acquisition of ectopic expression occurred in the lineage after the divergence of *D. guttifera* from its closely related species [17]. Thus, the co-option of *wg* that leads to the expression at the future pigmentation sites was one of the key events in the gain of the polka-dotted pigmentation pattern in *D. guttifera*.

**Fig. 1.**
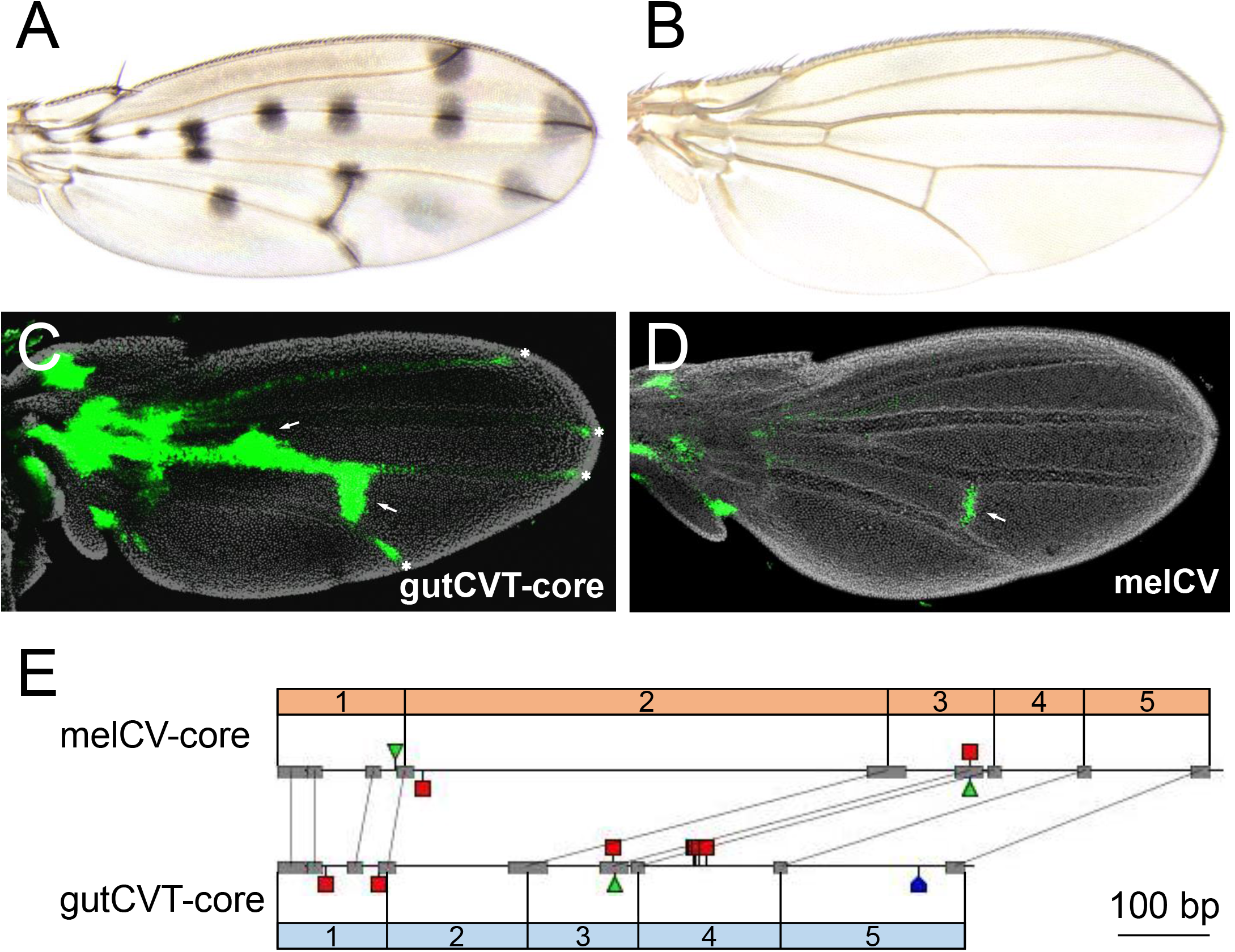
Adult wing of *D. guttifera* and *D. melanogaster* and *cis*-regulation of *wingless*. (A) Adult wing of *D. guttifera*. (B) Adult wing of *D. melanogaster*. (C) *wg* expression pattern driven by gutCVT-core. Reporter EGFP is expressed in the crossveins (arrows), and the longitudinal vein tips (asterisks). (D) *wg* expression pattern driven by melCV Reporter EGFP is expressed in the crossvein only. (E) Comparison of gutCVT-core and melCV-core. Gray bars indicate sequences that match at least 14 bp, visualized as anchor-points in GenePalette [38] and the two *cis*-regulatory sequences are divided into five regions by some of these anchor-points. Each divided region is called region 1, 2, 3, 4, or 5. Red squares indicate the sequences similar to SMAD consensus (GCCGnCGC or 1 bp change of it), blue pentagon indicates the sequence similar to BMP-AE (GCTGTGTAACCGGGG), and green triangles indicate the Medea binding motif (GTCTG). A–D were reproduced from [17].

A 760 bp *cis*-regulatory region in the vicinity of *wg* that drives expression in the crossveins and longitudinal vein tips (gutCVT-core) was identified in *D. guttifera* (Fig. 1C, E). The corresponding region of *D. melanogaster* (melCV-core and its larger version melCV) drives expression only in the crossveins (Fig. 1D, E). gutCVT-core maintains its function to regulate gene expression when introduced into *D. melanogaster*, and thus the new expression pattern of *wg* should have evolved via the changes in this *cis*-regulatory region rather than via changes in the *trans*-regulatory factors (Fig. 1C) [17, 20]. Then, what specific changes in the sequence were necessary to gain the new expression pattern?

In this research, we altered the sequences of the cloned *cis*-regulatory regions and examined their regulatory function by transgenic EGFP reporter assays. By deleting a part of gutCVT-core, we identified regions essential for the regulatory function and show that they are divided into regions needed to activate expression in the entire wing veins and a region required for repressing expression in excess parts of wing veins. In addition, by investigating the regulatory function of chimeric sequences with the homologous sequence of *D. melanogaster*, we revealed that some of the regions required for the regulation of the expression possessed their functions before the *cis*-regulatory evolution. Knockout of the potential binding sites of transcription factors revealed that a sequence resembling the SMAD binding site is necessary for activating *wg* expression and that this binding site also exists in the homologous region of *D. melanogaster*. These results suggest that the functions of pre-existing sequences and newly gained sequences were combined to acquire a new expression pattern of *wg*.

## Results

### Identification of the regions required for activating and repressing expression

The newly gained sequences that brought about the new expression pattern of *wg* should be included in sequences required for regulating expression. Thus, we first narrowed down where the sequences necessary for the regulatory function are located. gutCVT-core and melCV-core have several perfectly matched stretches of sequence (anchor-points in GenePalette function, 14–43 bp), and we divided these *cis*-regulatory regions into five regions bordered by anchor-points (Fig. 1E). *Cis*-regulatory sequences that lack one or some of these regions were then cloned next to an EGFP reporter gene. By integrating these constructs into a specific site in the genome of *D. melanogaster*, we examined their regulatory function in developing pupal wings (Fig. 2A). While complete gutCVT-core drove reporter EGFP expression in crossveins and longitudinal vein tips in pupal wings, gut2-5, which lacks region 1, activated expression in entire wing veins (Fig. 1C, 2B). gut3-5, lacking both regions 1 and 2, did not drive expression (Fig. 2C). These results indicate that gutCVT-core requires region 1 for repressing expression and region 2 for activating expression. gut1-4, which lacks region 5, drove EGFP expression in the crossveins and vein tips, which is very similar to the complete gutCVT-core (Fig. 2D). gut1-3, lacking both regions 4 and 5, activated expression only in the crossveins, and the EGFP expression pattern was similar to the one driven by melCV-core, but the fluorescence appeared weak (Fig. 2E). These results suggest that region 4 has an important role in activating expression, while region 5 is not necessary. Moreover, gut2-3, which lacks regions 1, 4, and 5, did not drive expression (Fig. 2F). This result indicates that region 1 has a role to activate expression in the crossveins.

**Fig. 2.**
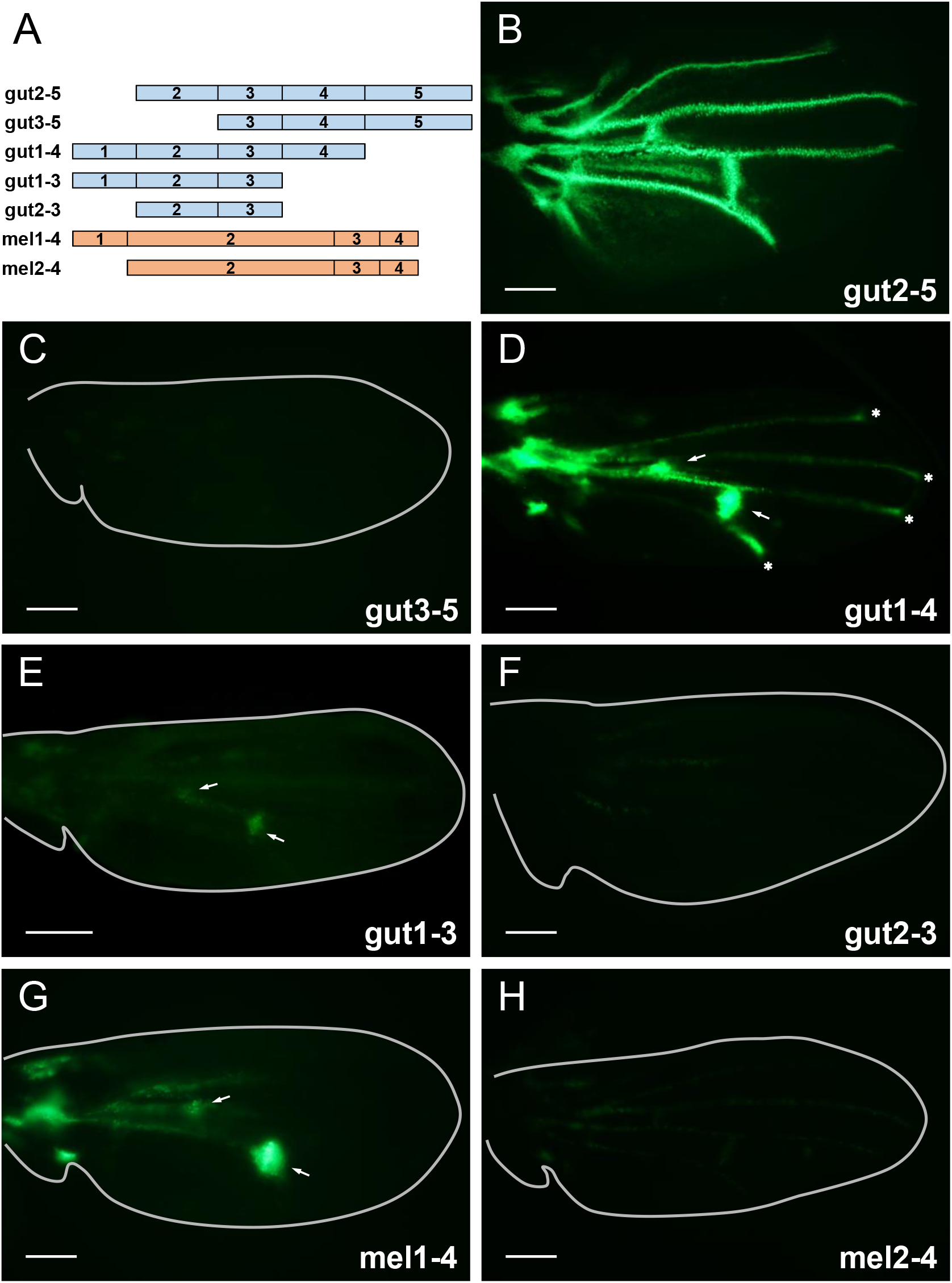
EGFP reporter assay for *cis*-regulatory sequences lacking some of the regions. (A) Enhancer activity of partial sequences of gutCVT-core (blue) and melCV-core (orange) was examined. (B) gut2-5 drove the reporter expression in the entire wing veins. (C) gut3-5 did not show strong expression in the wing veins. (D) gut1-4 activated expression in the crossveins (arrows) and longitudinal vein tips (asterisks). (E) gut1-3 drove expression in the crossveins (arrows) only. (F) gut2-3 did not show strong expression in the wing veins. (G) mel1-4 drove strong expression in the crossveins (arrow). (H) mel2-4 did not drive expression in the wing veins. White bars indicate 100 μm (B-H).

Similarly, to examine the regulatory function of the corresponding regions in melCV-core, we tested mel1-4, which lacks region 5, and mel2-4, which lacks regions 1 and 5. Similar to the complete melCV-core, mel1-4 activated expression in the crossveins (Fig. 2G). Thus, region 5 was also unnecessary for the function of melCV-core. In contrast, mel2-4 failed to activate the expression (Fig. 2H). Thus, it was shown that region 1 has an important role in driving expression in melCV-core while region 2–4 alone cannot activate expression in veins.

### Comparison of the regulatory function with a homologous sequence

The function to activate expression was not harbored in the same way in gutCVT-core and melCV-core. The activation function of region 2–4 was only possessed by gutCVT-core: then is the repressive function of region 1 only possessed by gutCVT-core? To test whether the repressive function of region 1 was newly gained in the lineage leading to *D. guttifera*, we examined the regulatory function of mel1gut2-5, a chimeric sequence in which the region 1 of gutCVT-core is replaced with the corresponding sequence of melCV-core. mel1gut2-5 showed a fluorescence pattern similar to gutCVT-core, with expression repressed in longitudinal veins except for the vein tips (Fig. 3A). Thus, the homologous sequence of *D. melanogaster* also has the repressive function of region 1.

**Fig. 3.**
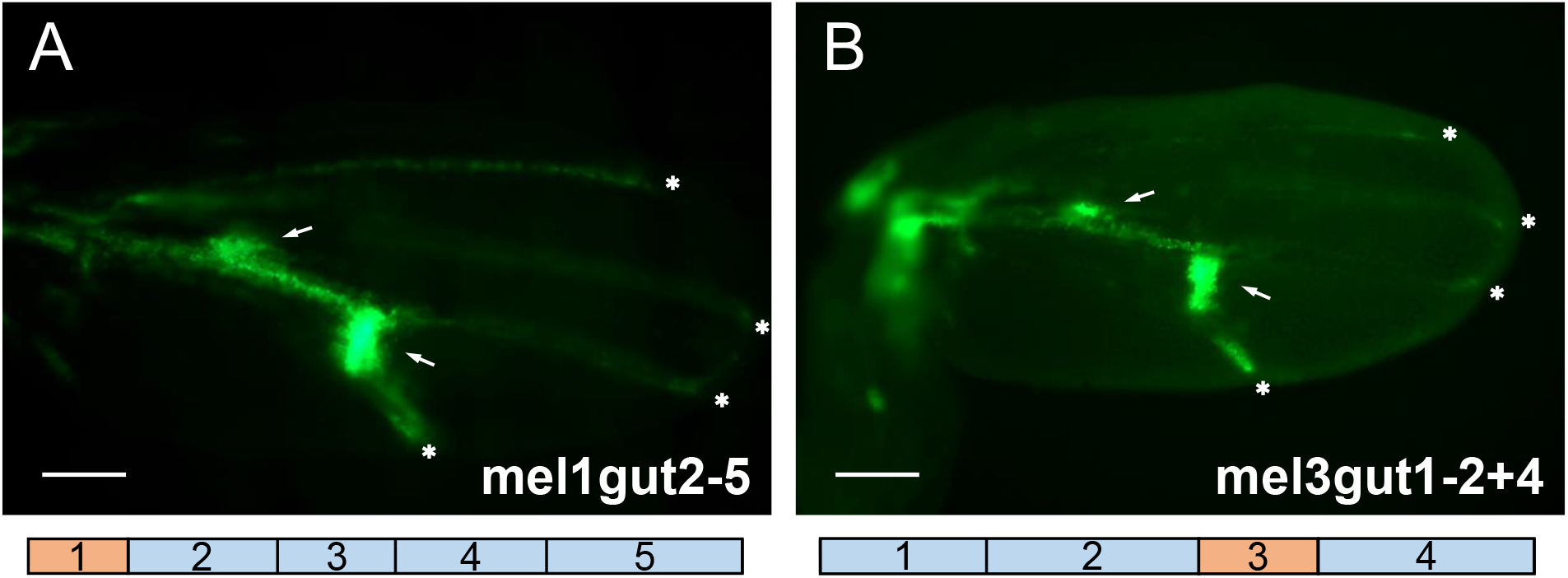
EGFP reporter assay for chimeric sequences. (A) Chimeric sequence in which region 1 of gutCVT-core is replaced with that of melCV-core. mel1gut2-5 drove reporter EGFP expression in the crossveins (arrows) and longitudinal vein tips (asterisks). (B) Chimeric sequence in which sequences from gutCVT-core are used for regions 1, 2, and 4, and a sequence from melCV-core is used for region 3. mel3gut1-2+4 activates expression in the crossveins (arrows) and longitudinal vein tips (asterisks). White bars indicate 100 μm (A, B).

In addition, since we could not confirm the necessity of region 3 in the previous experiment, we also tested the function of mel3gut1-2+4, a chimeric sequence in which region 3 is replaced with the corresponding sequence of melCV-core. This *cis*-regulatory sequence also has a regulatory function similar to gutCVT-core, where EGFP was expressed in the crossveins and longitudinal vein tips (Fig. 3B). This result indicates that changes in region 3 did not play a critical role in the *cis*-regulatory evolution that brought about the new expression pattern.

### Knockout of SMAD binding sites

Since the most important sequences for the regulatory function of a *cis*-regulatory region are binding sites of transcription factors, we focused on transcription factors expressed in the pupal wing. One of them, Mothers against Decapentaplegic (Mad), is a transcription factor required for Decapentaplegic signaling readout, and phosphorylated Mad (pMad) forms a complex with Medea, also a transcription factor, and binds to specific sequences to activate target gene expression [21–25]. The localization of pMad in the pupal wings is similar to the pattern driven by gutCVT-core [26, 27].

We hypothesized that Mad is the transcription factor that binds to gutCVT-core. We searched for candidate binding sites of Mad-Medea complex, BMP-AE (GGCGCCANNNNGNCV) [28], and candidate binding sites of Mad, SMAD consensus (GCCGNCGC) [21], in the *cis*-regulatory regions. In gutCVT-core, there are these binding sequences or very similar sequences, and one sequence similar to BMP-AE (GGGGCCAATGTGTCG) (hereinafter called BMP-AE for simplicity) was in region 5, two sequences similar to SMAD consensus (GCCGNCGC or 1 bp change of it) (hereinafter called SMAD consensus for simplicity) were in region 1 (CCCGCCGC and GCCGCCAC), one in region 3 (GCAGACGC), and four in region 4 (GCAGCCGC, GCCGCCGC, GCCGCCTC and GCCGCCTC) with overlap (Fig. 1E, 4A). To test the necessity of these putative binding sites for regulating expression, we knocked out these sequences and examined their regulatory function (Fig. 4A). As predicted from our result with gut1-4, BMP-AE KO possessed regulatory function very similar to that of gutCVT-core (Fig. 4B). On the other hand, gutSMAD bs KO, lacking all SMAD consensuses, failed to activate expression in the crossveins and vein tips, and EGFP was weakly expressed only around the crossveins and a part of the longitudinal vein between two crossveins (a region where *wg* expression was not shown by *in situ* hybridization) (Fig. 4C). This demonstrates that SMAD consensus is essential for activating expression. To identify which of the multiple SMAD consensuses is necessary for activating expression, we recovered SMAD consensuses in regions 3 and 4, which can be involved in promoting expression, respectively, and tested their regulatory function. gutSMAD bs3 Rec, in which only SMAD consensus in region 3 was recovered, regained its function to promote expression and drove EGFP expression in the crossveins and longitudinal vein tips (Fig. 4D). On the other hand, gutSMAD bs4 Rec, in which only SMAD consensuses in region 4 were recovered, failed to activate the expression and showed a non-specific EGFP expression pattern similar to gutSMAD bs KO (Fig. 4E). Thus, the only SMAD consensus required for regulating expression in gutCVT-core is the one in region 3.

**Fig. 4.**
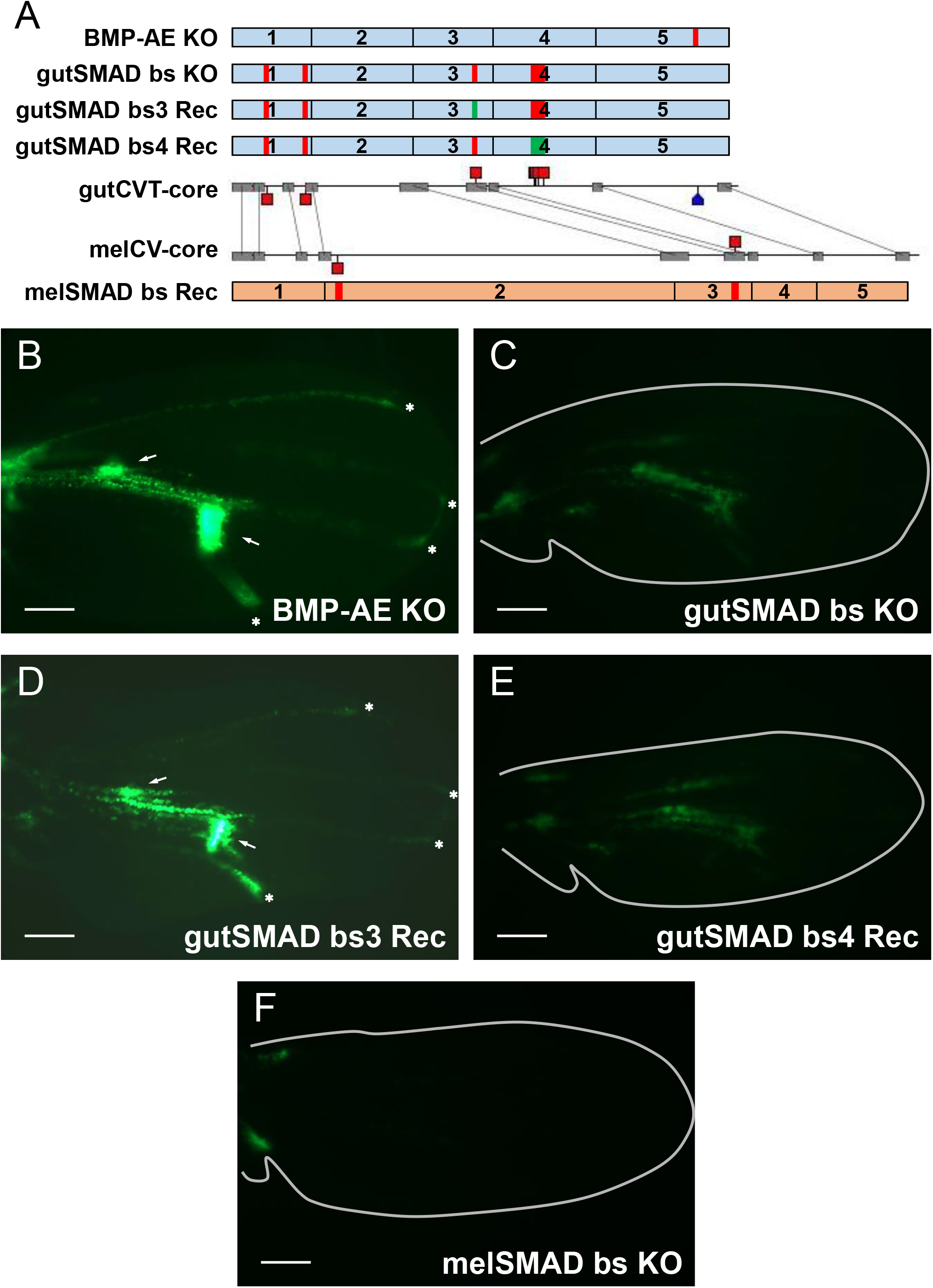
Changes in regulatory function by the knockout of candidate binding sites. (A) Sequences modified from gutCVT-core (blue) and melCV-core (orange) were examined. Red indicates the sections where the binding sites were knocked out and green indicates the sections where the knocked-out sequences were recovered (same as wildtype). (B) gutCVT-core whose BMP-AE was knocked out (BMP-AE KO) drove EGFP expression in crossveins (arrows) and longitudinal vein tips (asterisks). (C) gutCVT-core whose SMAD consensuses were all knocked out (gutSMAD bs KO) lost the enhancer activity in the wing veins. (D) When SMAD consensus in region 3 was recovered (gutSMAD bs3 Rec), the sequence regained the enhancer activity in the crossveins (arrows) and longitudinal vein tips (asterisks). (E) Even when all SMAD consensuses in region 4 were recovered (gutSMAD bs4 Rec), the sequence did not activate expression in the wing veins like gutSMAD bs KO. (F) When SMAD consensuses in melCV-core were all knocked out (melSMAD bs KO), the sequence lost the enhancer activity in the crossveins. White bars indicate 100 μm (B-F).

While no BMP-AE was found in melCV-core, two sequences very similar to SMAD consensus were present. One was in region 2 (GACGACGC), and the other was in region 3, which was completely identical to the sequence in gutCVT-core (GCAGACGC, Fig. 1E, 4A). To determine whether SMAD consensus is essential in melCV-core, we knocked out these sequences (melSMAD bs KO) (Fig. 4A). Deletion of SMAD consensus resulted in the loss of the EGFP expression in the crossveins, indicating that the SMAD consensus is required for melCV-core to activate expression (Fig. 4F).

## Discussion

### New expression pattern brought by the gain of enhancer activity in the entire wing veins

Although gutCVT-core expresses *wg* in the crossveins and longitudinal vein tips, this expression pattern was not simply formed by the activated expression in these parts of wing veins, but by a combination of activation in the entire wing veins by region 2–4 and repression in excess parts of longitudinal veins by region 1 (Fig. 5).

**Fig. 5.**
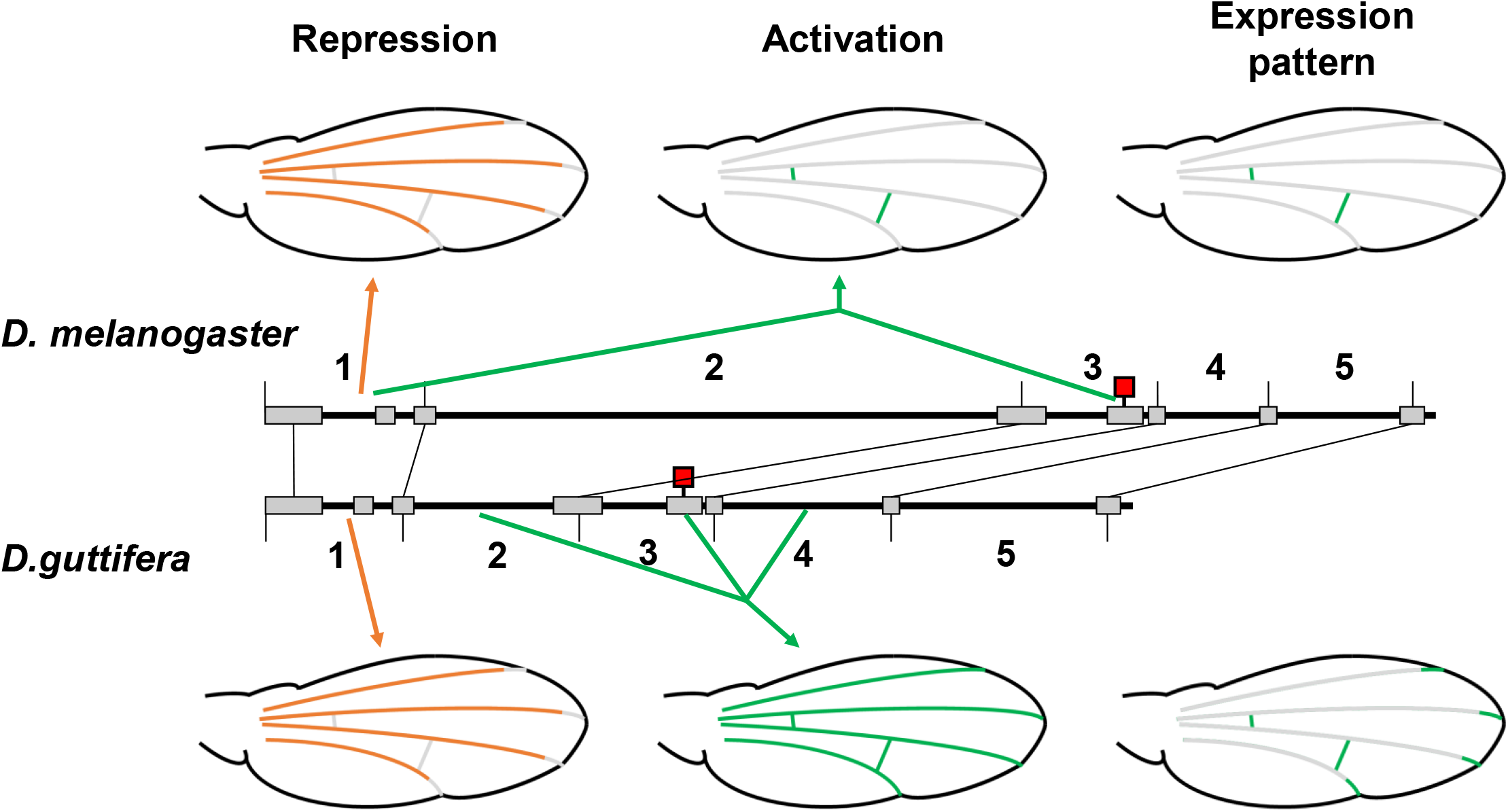
Predicted evolution of the sequence and the function of gutCVT-core. Repression of the gene expression in the longitudinal veins except the vein tips was possessed by both melCV-core and gutCVT-core (left, orange). In melCV-core, expression was activated only in the crossveins by sequences in region 1 and SMAD consensus in region 3 (red square) (upper center, green). In gutCVT-core, sequences that coordinate with the SMAD consensus in region 3 were acquired, producing a set of sequences which has activating function in the entire wing veins (lower center, green). In gutCVT-core, gene expression was driven in crossveins and longitudinal vein tips by the combination of the ancestral repression and the newly gained activation in the entire wing veins (lower right).

In addition, the repressing function of region 1 of gutCVT-core to repress expression throughout the longitudinal veins except the vein tips was also present in the homologous region of melCV-core, suggesting a silencer present in both species (Fig. 2B, 3A). Activating function of expression in the entire wing veins by region 2-4 was not possessed by melCV-core, and was shown to be newly gained in gutCVT-core (Fig. 2B, H). Thus, it was revealed that the new expression of *wg* in the longitudinal vein tips in *D. guttifera* was not simply brought by the gain of activation in these parts, but rather the activation in the entire wings.

One interesting point is that melCV-core possesses the repressive function in region 1, even though the region 2–4 has no function to activate gene expression in the entire wing veins. The role of the sequence involved in repressing expression in melCV-core is still unknown, but one possibility is that this sequence originally works in developmental stages other than pupae or in other tissues. *Cis*-regulatory sequences sometimes change the place where or the developmental timing when they work over the course of evolution [29]. Alternatively, an ancestor of *D. melanogaster* might have had activating function in the region 2–4 and even pigmentation on veins like *D. guttifera*, and lost them secondarily, although this scenario requires multiple assumptions and we consider it unlikely.

### The role of the pre-existing SMAD consensus in the gain of the new regulatory function of *wg* expression

SMAD consensus in region 3 was required for activating expression in the entire wing veins (Fig. 4). However, this does not mean that the gain of the SMAD consensus brought about the ectopic expression of *wg*. The SMAD consensus required for promoting expression is also conserved in melCV-core, and the chimeric sequence in which region 3 was replaced with a homologous sequence of melCV-core drove the same expression pattern as full gutCVT-core (Fig. 3B). Furthermore, also in melCV-core, the deletion of all SMAD consensus causes the loss of gene expression in the crossveins, suggesting that the SMAD consensus in region 3, which is conserved between the two species, is necessary for activating expression by melCV-core (Fig. 4F). Thus, we propose that the SMAD consensus in ancestral *cis*-regulatory regions coordinates with some newly gained sequences to acquire *wg* expression in the new parts of the wing veins (Fig. 5).

The causal sequence change in gutCVT-core responsible for the evolution of the *wg* expression pattern has not yet been identified at nucleotide level resolution. That sequence is presumed to coordinate with SMAD consensus to activate expression in the entire wing veins. Therefore, newly gained elements may interact with SMAD proteins. For example, when pMad works to activate transcription, it makes a complex with Medea and the complex binds to BMP-AE, but the SMAD consensus is shorter than BMP-AE and does not contain the binding sites of both [21–25]. There was one Medea consensus on the SMAD consensus in region 3, but they are unlikely to work together because the binding consensuses were overlapped. Medea consensus is short (GTCTG, [30]) and similar sequences are abundant within gutCVT-core; thus, Medea may bind to some of them. Some factors are known to interact with pMad. For example, the protein CBP/p300 works as a transcriptional co-activator [31, 32]. Such Mad-related factors may provide clues for identifying newly gained regulatory sequences in gutCVT-core.

### Acquisition of the new expression pattern by the combination of pre-existing and newly gained *cis*-elements

In the evolution of *wg* expression pattern associated with the wing pigmentation pattern formation of *D. guttifera*, at least two pre-existing *cis*-regulatory sequences were co-opted. One is the sequence in region 1 to repress expression throughout the longitudinal veins except for the vein tips, and the other is the SMAD consensus in region 3 required for activating expression in the entire wing veins. These pre-existing *cis*-regulatory sequences function together with newly gained *cis*-regulatory sequences in gutCVT-core, resulting in the new expression pattern of *wg* (Fig. 5). This means that not all of the sequences required for the new expression pattern were newly gained in the co-option of *wg*. Similar to this case, there are several examples of new expression patterns evolved by the modification of pre-existing enhancers [33–35].

Combination of multiple control functions could possibly produce diverse expression patterns. Such *cis*-regulatory regions would require many regulatory sequences, and therefore the probability of acquiring a new one from a non-functional sequence by mutation seems intuitively low. The frequency of *de novo* generation of enhancers is still being discussed [36]. Alternatively, by reusing preexisting regulatory sequences, fewer sequences are required to gain new expression patterns. Co-option of pre-existing sequences is one of the main mechanisms that cause the gain of ectopic expression of a gene, and if it occurs on important regulatory genes, as shown in this study, may lead to gene network co-option [37].

## Materials and Methods

### Prediction of binding sites in *cis*-regulatory regions

To identify putative binding sites, we used Gene Palette 2.1 [38]. To find candidates for BMP-AE, we searched for GGCGCCANNNNGNCV [28], allowing a 1 bp mismatch, in gutCVT-core and melCV-core. In the same way, we found the putative SMAD consensus by searching for GCCGNCGC [21], allowing a 1bp mismatch, in the two *cis*-regulatory regions.

### Examination of the function of modified *cis*-regulatory sequences by EGFP reporter assay

We first amplified *cis*-regulatory sequences to test the regulatory function by PCR, using gutCVT-core or melCV-core [17] as templates. To synthesize sequences that lack some of the regions, PCR was conducted using primers based on sequences from the borders of the divided regions. To obtain chimeric sequences, we made DNA fragments of the respective regions of gutCVT-core or melCV-core and joined them together by overlap extension PCR. To obtain sequences in which putative binding sites are knocked out, we first performed PCR using primers that contain sequences of about 10 bases before and after the sequence to be knocked out, with the putative binding sites changed to random sequences (in the case of BMP-AE KO, all bases were changed to A). These were then joined together by overlap extension PCR. By using PROMO [39, 40], we confirmed that no important binding site was newly created. All primers used in this research are shown in Supporting Information (Table S1, S2).

Amplified sequences were then inserted into S3aG [41] with the 5’ end oriented closer to the EGFP sequence and the constructs were then introduced into *D. melanogaster*, VK00006 (attP on cytogenetic location 19E7) [42] using PhiC31 integrase-mediated transgenesis systems (BestGene, CA). Transgenic flies were then crossed to be homozygous for the introduced sequence and maintained at 25 °C with standard cornmeal/sugar/yeast/agar food [43].

To observe EGFP reporter expression, wandering larvae were collected and then incubated at 25 °C for 40–44 hours, and we obtained P6-P7 pupa [44]. In these stages, pupal wings have not yet folded, and wing veins are visible. Obtained pupas were taken out from the puparium, had their fat and organs removed in 1 x PBS, and were fixed in 1% paraformaldehyde at 4 °C overnight. Fixed wings were taken out from pupal cuticles in 1 x PBS and observed in the fluorescence field using a BX60 microscope (Olympus) [43]. To examine the EGFP expression pattern on wings, we took photographs under the same conditions of light intensity and exposure. The brightness and contrast of images were adjusted with ImageJ [45].

Fluorescence patterns were confirmed in one line each for BMP-AE KO, gut1-3, and gutSMAD bs3 Rec, and in multiple lines for each of the other constructs.

## Abbreviations

EGFP: Enhanced Green Fluorescent Protein
Mad: Mothers against Decapentaplegic
BMP-AE: Bone Morphogenetic Protein-Activating Element

## Author contributions

TK, NS, and SK conceived and designed the study. TK and NS collected data and conducted the analysis. TK and SK wrote the manuscript. All authors gave final approval for publication and agreed to be held accountable for the work performed therein.

## Acknowledgements

We thank Sean B. Carroll for providing template DNA; Elizabeth Nakajima for English editing; Masato Koseki for technical advice; and Haruka Takahashi, Masato Koseki, Satoshi Hayakawa, Shunsuke Otake, Sota Sakai, Takuma Niida, Yusuke Yoshida and Yuto Terashima for their discussions and comments. This work was funded by KAKENHI (18H02486) to SK.

