## Supplementary material for "*Cis*-regulatory evolution that caused change in *wingless* expression pattern associated with wing pigmentation pattern of *Drosophila*": Table S1, S2

Table. S1. All primers used in this research

| Primer name | Pimer Sequence | Resriction Site |
|---|---|---|
| gut BMP-AE knockout 1 AscI R1 | TCTTGGCGCGCCCTCGAGCACAACAATTGCCACTTAATTAACAC GAACTCCAACTTCTTTTTTTTTTTTTTTAAAATACATATTTCG | AscI |
| gut JY1A SMAD knockout1 SbfI F1 | TAATCCTGCAGGGACATTGCTCCTAATCAATAAAACTAACGAGC GGATTAAGATTATGTTGCACGATAACTCATCCACACAGAGGCG | SbfI |
| gut JY1L SMAD knockout1 F1 | GTGTTGTAATTTGACGTCCCACACGTTTGC | |
| gut JY1L SMAD knockout1 R1 | GCAAACGTGTGGGACGTCAAATTACAACAC | |
| gut JY1M SMAD knockout1 F1 | CAAAAATGGTTTGTCATCGCTAATGAATTGC | |
| gut JY1M SMAD knockout1 R1 | GCAATTCATTAGCGATGACAAACCATTTTTG | |
| gut multiSMAD knockout1 F1 | AGACATTGAGAGCGCAACGCGTATTGTTACACGTCGCCTTCT | |
| gut multiSMAD knockout1 R1 | AGAAGGCGACGTGTAACAATACGCGTTGCGCTCTCAATGTCT | |
| gut wg JY1J AscI R1 | TCTTGGCGCGCCCTCGAGCACAACAA | AscI |
| gut wg JY1L AscI R1 | ATATGGCGCGCCCCAAATCAAACAGCGGCGCA | AscI |
| gut wg JY1L F1 Karasawa | CGCTGTTTGATTTGGCCAAA | |
| mel JY1L SMAD knockout1 F1 | TGTTGTAATTTGCGAACCCACACGTTTCG | |
| mel JY1L SMAD knockout1 R1 | CGAAACGTGTGGGTTCGCAAATTACAACA | |
| mel JY1M SMAD knockout1 F1 | CGAGTCGCCATCATGCATCACTATAAG | |
| mel JY1M SMAD knockout1 R1 | CTTATAGTGATGCATGATGGCGACTCG | |
| mel wg JY1J AscI R1 | TCTTGGCGCGCCCTCACGCCTCGAAAC | AscI |
| mel wg JY1L R1 Karasawa | CCAAATCAAACAGCGCGCGA | |
| wg JY1A SbfI F2 | ATCGCCTGCAGGGACATTGCTCCTAA | SbfI |
| wg JY1M For Karasawa | TGGCGGCCTAATGAATTGC | |
| wg JY1M Rev Karasawa | GCAATTCATTAGGCCGCCA | |
| wg JY1M SbfI F1 | TAATCCTGCAGGTGGCGGCCTAATGAATTGC | SbfI |
| wg JY1O AscI R1 | ATATGGCGCGCCTTGTCATGCCAAGAGTGGCG | AscI |
| wg JY1P F1 Karasawa | GATGTGACAATGACAGTTGTTGTTC | |
| wg JY1P R1 Karasawa | GAACAACAACTGTCATTGTCACATC | |
| wg JY1P SbfI F1 | TAATCCTGCAGGGATGTGACAATGACAGTTGTTGTTC | SbfI |

Table. S2. Primers used for each tested *cis*-regulatory sequences

| Tested sequence | Primers for producing dna fragments | Template sequence |
|---|---|---|
| gut2-5 | wg JY1M SbfI F1 | gutCVT-core |
| | gut wg JY1J AscI R1 | gutCVT-core |
| gut3-5 | wg JY1P SbfI F1 | gutCVT-core |
| | gut wg JY1J AscI R1 | gutCVT-core |
| gut1-4 | wg JY1A SbfI F2 | gutCVT-core |
| | wg JY1O AscI R1 | gutCVT-core |
| gut1-3 | wg JY1A SbfI F2 | gutCVT-core |
| | gut wg JY1L AscI R1 | gutCVT-core |
| gut2-3 | wg JY1M SbfI F1 | gutCVT-core |
| | gut wg JY1L AscI R1 | gutCVT-core |
| mel1-4 | wg JY1A SbfI F2 | melCV-core |
| | wg JY1O AscI R1 | melCV-core |
| mel2-4 | wg JY1M SbfI F1 | melCV-core |
| | wg JY1O AscI R1 | melCV-core |
| mel1gut2-5 | wg JY1A SbfI F2 | melCV-core |
| | wg JY1M Rev Karasawa | melCV-core |
| | wg JY1M For Karasawa | gutCVT-core |
| | gut wg JY1J AscI R1 | gutCVT-core |
| | wg JY1A SbfI F2 | overlap extension PCR |
| | gut wg JY1J AscI R1 | overlap extension PCR |
| mel3gut1-2+4 | wg JY1A SbfI F2 | gutCVT-core |
| | wg JY1P R1 Karasawa | gutCVT-core |
| | wg JY1P F1 Karasawa | melCV-core |
| | mel wg JY1L R1 Karasawa | melCV-core |
| | gut wg JY1L F1 Karasawa | gutCVT-core |
| | wg JY1O AscI R1 | gutCVT-core |
| | wg JY1A SbfI F2 | overlap extension PCR |
| | wg JY1O AscI R1 | overlap extension PCR |
| BMP-AE KO | wg JY1A SbfI F2 | gutCVT-core |
| | gut BMP-AE knockout 1 AscI R1 | gutCVT-core |
| gutSMAD bs KO | gut JY1A SMAD knockout1 Sbf1 F1 | gutCVT-core |
| | gut JY1M SMAD knockout1 R1 | gutCVT-core |
| | gut JY1M SMAD knockout1 F1 | gutCVT-core |
| | gut JY1L SMAD knockout1 R1 | gutCVT-core |
| | gut JY1L SMAD knockout1 F1 | gutCVT-core |
| | gut multiSMAD knockout1 R1 | gutCVT-core |
| | gut multiSMAD knockout1 F1 | gutCVT-core |

| | | |
|---|---|---|
| | gut wg JY1J AscI R1 | gutCVT-core |
| | wg JY1A SbfI F2 | overlap extension PCR |
| | gut wg JY1J AscI R1 | overlap extension PCR |
| gutSMAD bs3 Rec | gut JY1A SMAD knockout1 Sbf1 F1 | gutCVT-core |
| | gut JY1M SMAD knockout1 R1 | gutCVT-core |
| | gut JY1M SMAD knockout1 F1 | gutCVT-core |
| | gut multiSMAD knockout1 R1 | gutCVT-core |
| | gut multiSMAD knockout1 F1 | gutCVT-core |
| | gut wg JY1J AscI R1 | gutCVT-core |
| | gut JY1A SMAD knockout1 Sbf1 F1 | overlap extension PCR |
| | gut wg JY1J AscI R1 | overlap extension PCR |
| gutSMAD bs4 Rec | gut JY1A SMAD knockout1 Sbf1 F1 | gutCVT-core |
| | gut JY1M SMAD knockout1 R1 | gutCVT-core |
| | gut JY1M SMAD knockout1 F1 | gutCVT-core |
| | gut JY1L SMAD knockout1 R1 | gutCVT-core |
| | gut JY1L SMAD knockout1 F1 | gutCVT-core |
| | gut wg JY1J AscI R1 | gutCVT-core |
| | gut JY1A SMAD knockout1 Sbf1 F1 | overlap extension PCR |
| | gut wg JY1J AscI R1 | overlap extension PCR |
| melSMAD KO | wg JY1A SbfI F2 | melCV-core |
| | mel JY1M SMAD knockout1 R1 | melCV-core |
| | mel JY1M SMAD knockout1 F1 | melCV-core |
| | mel JY1L SMAD knockout1 R1 | melCV-core |
| | mel JY1L SMAD knockout1 F1 | melCV-core |
| | mel wg JY1J Asc1 R1 | melCV-core |
| | wg JY1A SbfI F2 | overlap extension PCR |
| | mel wg JY1J Asc1 R1 | overlap extension PCR |
